# Ribosomal protein S7 ubiquitination during ER stress in yeast is associated with selective mRNA translation and stress outcome

**DOI:** 10.1101/2020.09.24.311365

**Authors:** Yasuko Matsuki, Yoshitaka Matsuo, Yu Nakano, Shintaro Iwasaki, Hideyuki Yoko, Tsuyoshi Udagawa, Sihan Li, Yasushi Saeki, Tohru Yoshihisa, Keiji Tanaka, Nicholas T. Ingolia, Toshifumi Inada

## Abstract

eIF2α phosphorylation-mediated translational regulation is crucial for global translation repression by various stresses, including the unfolded protein response (UPR). However, translational control during UPR has not been demonstrated in yeast. This study investigated ribosome ubiquitination-mediated translational controls during UPR. Tunicamycin-induced ER stress enhanced the levels of ubiquitination of the ribosomal proteins uS10, uS3 and eS7. Not4-mediated monoubiquitination of eS7A was required for resistance to tunicamycin, whereas E3 ligase Hel2-mediated ubiquitination of uS10 was not. Ribosome profiling showed that the monoubiquitination of eS7A was crucial for translational regulation, including the upregulation of the spliced form of HAC1 (*HAC1i*) mRNA and the downregulation of Histidine triad NucleoTide-binding 1 (*HNT1*) mRNA. Downregulation of the deubiquitinating enzyme complex Upb3-Bre5 increased the levels of ubiquitinated eS7A during UPR in an Ire1-independent manner. These findings suggest that the monoubiquitination of ribosomal protein eS7A plays a crucial role in translational controls during the ER stress response in yeast.

## INTRODUCTION

The protein folding capacity of cells is important for maintaining endoplasmic reticulum (ER) homeostasis. Accumulation of unfolded proteins in the ER induces the unfolded protein response (UPR), which consists of a set of signalling pathways that respond to the resulting ER stress^1^. In metazoan cells, the UPR is divided into three branches: inositol-requiring enzyme 1 (IRE1), ATF6 and PKR-like ER kinase (PERK)^1^. The IRE1 branch increases protein folding capacity by inducing translation of the transcription factor XBP1^2,3^; the ATF6 branch is involved in the increase of ER folding ability^4^; and the PERK branch reduces the initiation of global translation through the phosphorylation of eIF2α^5^. eIF2α phosphorylation, in turn, leads to the selective translation of transcription factors such as ATF4, thereby increasing ER folding ability^6^. A unique ISR/PERK-mediated translational control mechanism, independent of the eIF2α phosphorylation*/*eIF2B axis, is crucial for recovery from the chronic ER stress response^7^. In budding yeast, activated Ire1 on the ER removes non-conventional introns from unspliced *HAC1* mRNA (*HAC1*^u^)^8-12^, thereby relieving translational repression^13,14^. Translation of the spliced (induced) form of *HAC1* mRNA (*HAC1*^i^) produces Hac1, a transcription factor that induces the expression of UPR target genes, which play crucial roles in increasing the protein folding and degrading abilities of the ER^15^. Although the UPR is thought to induce translational regulation in the pathogenic fungus *Aspergillus fumigatus*^16^, translational control in response to the UPR has not been evident in the yeast *Saccharomyces cerevisiae*.

Specific modification of ribosomal proteins is thought to be critical for regulating translation in eukaryotes. One example is the role of ribosome ubiquitination in the quality control system for aberrant translation. Ribosome-associated Quality Control (RQC) is a translation arrest-induced quality control pathway that leads to the co-translational degradation of the arrested products^17-21^. In the first step of RQC, abnormal stalling ribosomes are recognized, and specific residues in the stalled ribosomes are ubiquitinated. In yeast, the E3 ubiquitin ligase Hel2 ubiquitinates uS10 at K6 and K8 and plays a crucial role in RQC^22-24^. ZNF598, the mammalian homologue of Hel2, ubiquitinates the ribosomal proteins eS10 at K138/K139 and uS10 at K4/K8, thereby inducing translational arrest and RQC triggered by poly-lysine sequences in mRNA^25-27^. Collided ribosomes form a unique structural interface to induce E3 ligase Hel2/ZNF598-driven quality control pathways^24,28^. Moreover, sequential ubiquitination of the ribosomal protein uS3 triggers the degradation of nonfunctional 18S rRNA^29^.

Despite increased understanding of the roles of ribosome ubiquitination in quality control pathways, the physiological relevance of ribosome ubiquitination remains largely unknown. However, ribosome ubiquitination has been linked to cellular responses to stress. For example, K63 polyubiquitination may modulate oxidative stress responses, and ubiquitination of specific lysine residues of ribosomal proteins may contribute to stress responses^30,31^. Although more than 100 ubiquitination sites have been identified in ribosomes, the physiological relevance of ribosome ubiquitination remains unclear. Induction of the UPR in mammalian cells was shown to upregulate the expression of the ubiquitinated ribosomal proteins uS10, eS10 and uS3^32^. Nevertheless, the exact role of ribosome ubiquitination in the UPR remains unknown.

The present study found that ribosome ubiquitination is essential for gene regulation during the UPR in yeast. The E3 ligase Not4-mediated ubiquitination of eS7A was required for resistance to tunicamycin (Tm), an ER stress-inducing compound. Ribosome profiling showed that ubiquitination of eS7A was required for translational controls during the UPR. These findings indicate that Not4-mediated monoubiquitination of eS7A is essential for controlling the translation of specific mRNAs during the ER stress response, including through the upregulation of *HAC1*^*i*^ mRNA and the downregulation of *HNT1* mRNA.

## RESULTS

### Monoubiquitination of eS7A is required for translational regulation during the UPR in yeast

Ribosome ubiquitination increases significantly upon induction of the UPR in mammalian cells^32^. To assess the role of ribosome ubiquitination in the UPR in yeast, the levels of ubiquitinated ribosomal proteins were evaluated by affinity purification of ribosomes with FLAG-tagged Rpl25 from cells expressing N-terminal Myc-tagged Ubiquitin protein (Ub), as previously described^22^. The levels of the ubiquitinated ribosomal proteins uS10, uS3 and eS7A were substantially increased under UPR conditions (Fig. 1a). Not4 is an E3 ligase for the ribosomal proteins eS7A and eS7B in yeast^33,34^, and is involved in translation repression^35^. Cells expressing *not4*Δ mutant showed Tm-sensitive growth (Fig. 1b). In addition, the *eS7a-4KR* mutant, which involves lysine-to-arginine substitutions at all four Not4-specific ubiquitination sites, had the same level of Tm sensitivity as *not4*Δ mutant cells (Fig. 1b). Monoubiquitinated eS7A was not detected in *not4*Δ mutant cells (Fig. 1c), confirming that Not4 is responsible for the monoubiquitination of eS7A during the UPR. Because Hel2 was previously reported to form K63-linked polyubiquitin chains on Not4 monoubiquitinated eS7A and to play a crucial role in No-Go Decay (NGD)^24^, we assessed the involvement of Hel2 in the UPR. In contrast to Not4, the deletion of Hel2 (*hel2*Δ) did not affect cell growth in the presence of Tm (Fig. 1b). In addition, neither the uS10 nor the uS3 mutant of Hel2-target lysine residues (*uS10-K6/8R* and *uS3-K212R*) affected cell growth in the presence of Tm (Fig. 1b). The uS10-Ub and uS3-Ub signals were not detected in the *hel2*Δ deletion mutant, whereas the eS7-Ub signal was not detected in the *not4*Δ deletion mutant (Supplementary Figure 1), suggesting that eS7 ubiquitination is dependent on Not4. We previously reported that Hel2 elongates the ubiquitin chain at eS7 after Not4-dependent monoubiquitination, which is essential for mRNA quality control in NGD^36^. However, in contrast to *not4*Δ and the eS7-4KR mutant, the *hel2*Δ mutant did not show sensitivity to Tm (Fig. 1b). These results suggest that Not4-mediated monoubiquitination of the ribosomal protein eS7A is indispensable for cell survival under ER stress conditions, whereas Hel2-mediated polyubiquitination of eS7A is not.

**Fig. 1.**
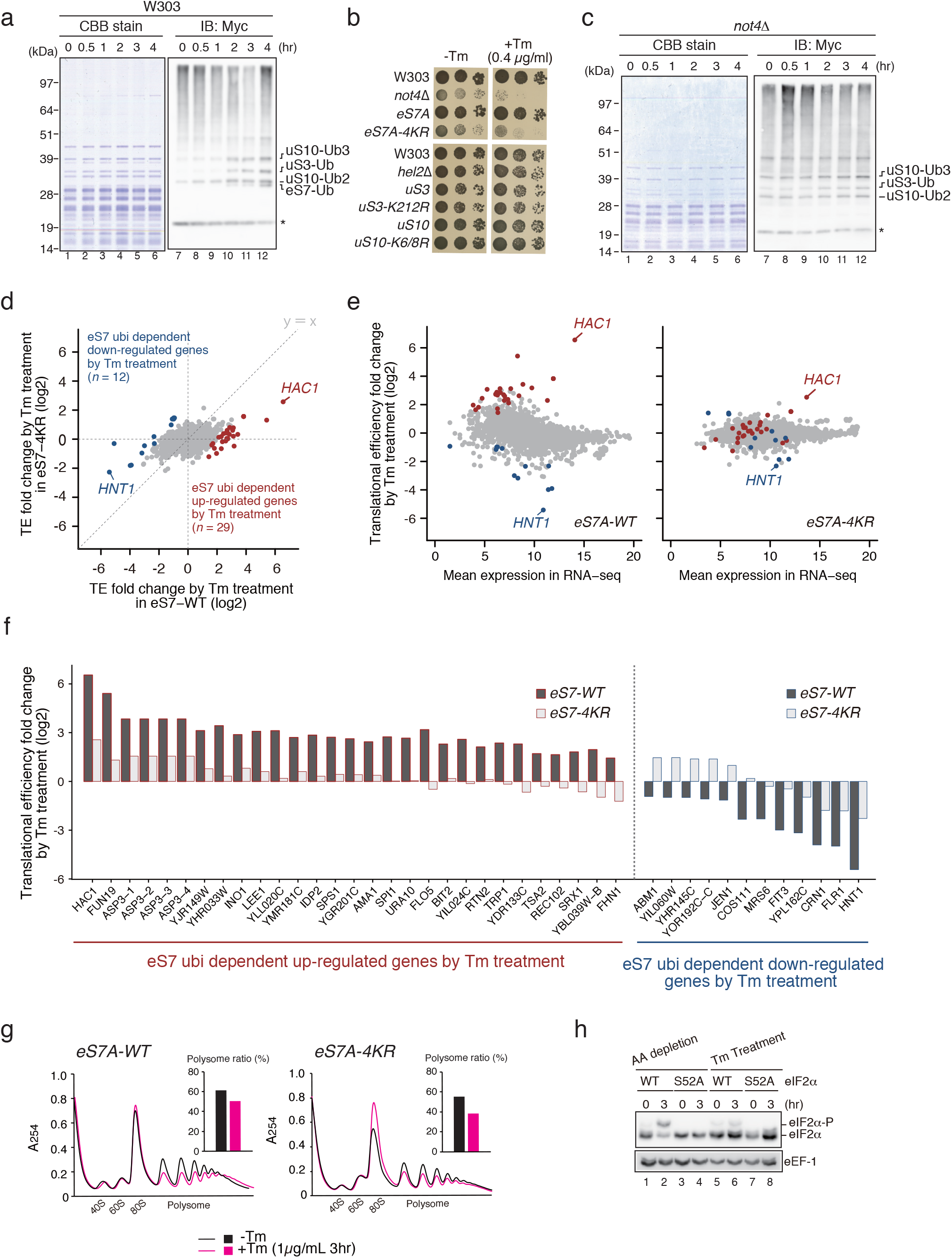
Not4-mediated monoubiquitinated eS7A is required for translational controls in the UPR in yeast. **a**, Ubiquitinated proteins in the ribosome after the addition of tunicamycin. Yeast cells harboring p*CUP1*p-*MYC-UBI* and p*RPS2(uS5)-FLAG* or p*RPL25(uL23)-FLAG* were cultured in 800 mL of synthetic complete medium. Myc-Ubi expression was induced by culturing the cells in the presence of 0.1 mM Cu^2+^ for 2 h. Cell lysates were prepared and FLAG-tagged ribosomes were purified using an M2 FLAG-affinity resin (Sigma), as described^56^. Affinity purified samples were subjected to SDS-PAGE followed by western blotting with an anti-Myc antibody. The arrows indicate proteins previously identified by mass spectrometry. **b**, The Not4 ubiquitination of ribosomal protein eS7A is crucial for UPR in yeast. Genetic screening was performed to identify the E3 ubiquitin enzyme NOT4 required for resistance to Tm. **c**, Dependence of eS7A mono- and poly-ubiquitination on Not4. **d**, Ribosome profiling showing up- and down-regulation of translation by the eS7A ubiquitination. The ribosome profiling and RNA-seq results represent two independent biological replicates. The correlations between replicates are shown in Supplementary Figure 3a-b. **f**, eS7A ubiquitination-dependent up- and down-regulation of specific mRNAs in response to UPR. The mRNA most upregulated by eS7A ubiquitination was *HAC1*, and the mRNA most upregulated was *HNT1*. **g**, UPR does not inhibit bulk translation in wild-type and mutant cells. **h**, Phosphorylation of eIF2α in response to amino acid depletion or Tm treatment. Shown are the levels of eIF2α phosphorylation in WT and S52A mutants in response to amino acid starvation and the presence of Tm. **a, c, h**, Cropped gels or blots were display. All uncropped images are available in Supplemental Figure. S7.

These results suggested that translational control may play a role in the UPR in yeast and may involve eS7A ubiquitination. To test this, we then performed RNA-seq and ribosome profiling to investigate the regulation of translation in response to ER stress, and estimated translation efficiency (TE) by assessing both mRNA abundance and ribosome occupancy. During the early response, within 1 h after Tm treatment, the translation of 20 mRNAs was statistically and significantly changed (Supplementary Fig. 2a; q-value <0.01), whereas >200 mRNAs were up- or down-regulated 4 h after Tm treatment (Supplementary Fig. 2a). eS7A ubiquitination-dependent translational regulation was therefore monitored at 4 h. A modest translational response was observed in *eS7A-4KR* mutant cells, with statistically significant changes in the translation of 214 mRNAs (Fig. 1d-e; Supplementary Fig. 2b; q-value <0.01). To examine the eS7A ubiquitination dependency of the involved mRNAs, TE was compared in *eS7A-WT* and *eS7A-4KR* mutant cells (Fig. 1d-e), with 29 and 12 mRNAs categorised as being up- and down-regulated, respectively, in response to eS7A ubiquitination (Fig. 1f). These subsets were identified using the formula “log2 TE fold change (*eS7A-WT*) -log2 TE fold change (*eS7A-4KR*)”, with mRNAs scored as >2 and <-2 defined being up- and down-regulated, respectively, in response to eS7A ubiquitination. Polysome analysis showed that UPR moderately repressed general translation in *wild-type* and *eS7A-4KR* mutant cells (Fig. 1g). The levels of phosphorylated eIF2α did not increase during the UPR (Fig. 1h), suggesting that initiation of general translation was not inhibited by phosphorylated eIF2α during the UPR. To rule out the effect of *eS7A-4KR* mutations on general translation, protein synthesis rates were measured in *eS7A-WT* and the *eS7A-4KR* mutant by puromycin labelling. Protein synthesis rates did not differ significantly in *eS7A-WT* and *eS7A-4KR* (Supplementary Fig. 3a-b), indicating that eS7A monoubiquitination is involved in regulating the translation of specific mRNAs in response to ER stress.

**Fig. 2.**
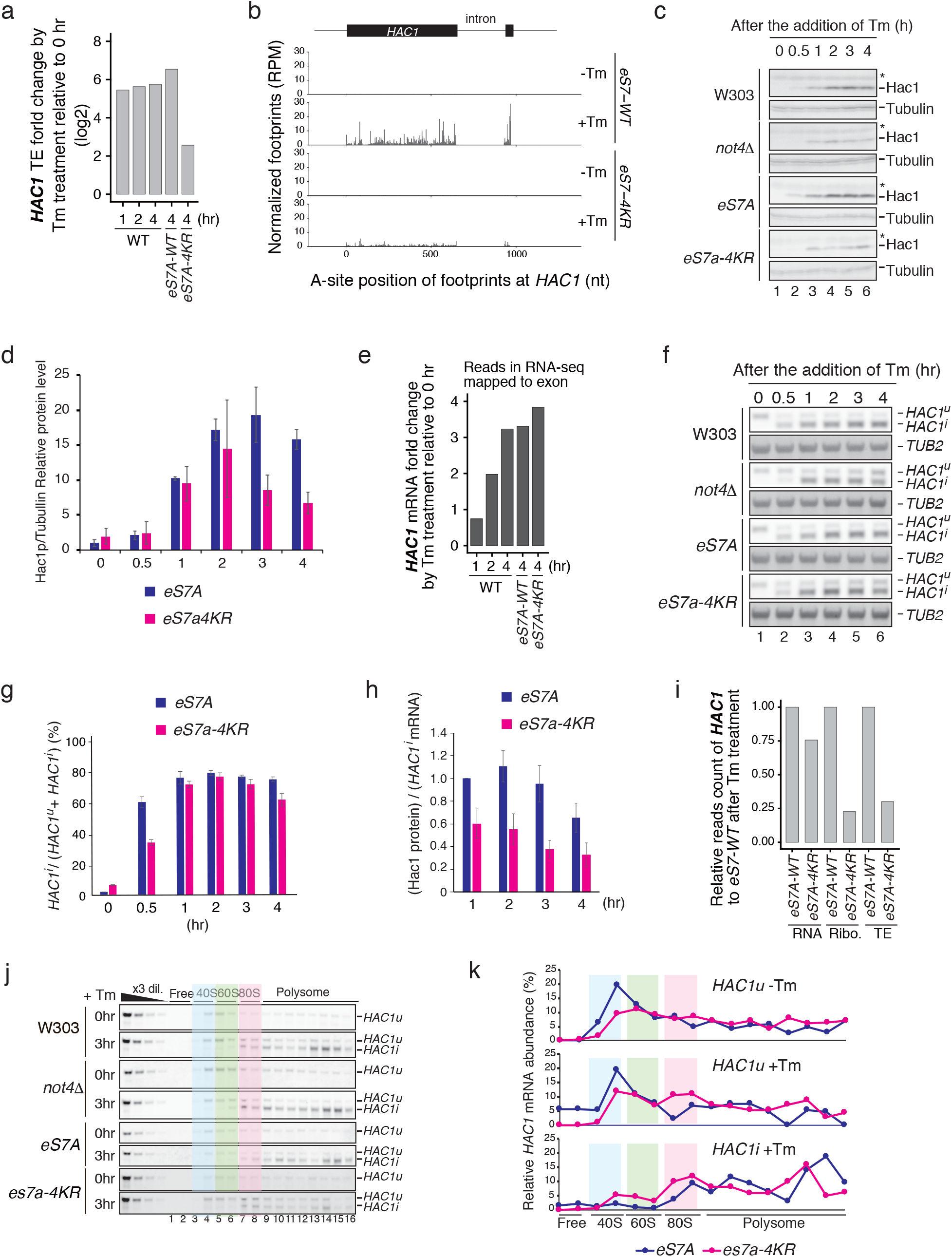
Not4-mediated monoubiquitinated eS7A is required for upregulation of *HAC1*^*i*^ translation in response to UPR in yeast. **a**, Upregulation of *HAC1*^*i*^ translation in response to UPR was diminished in *eS7A-4KR* mutant cells. **b**, Map of the A-site position of footprints at *HAC1*. **c**, Significant reductions in Hac1 protein levels in *not4*Δ and *eS7A-4KR* mutant cells. **d**, Normalisation of Hac1 protein levels relative to tubulin levels. The normalised levels of Hac1 protein were significantly lower in eS7A-4KR mutant than in WT. **e**, Upregulation of *HAC1* mRNA in W303(WT), *eS7A-WT* and *eS7A-4KR* mutant strains. **f**, Regulation of mRNA splicing of *HAC1*^*u*^ was moderately reduced in *eS7A-4KR* mutant cells. **g**, Splicing efficiency of *HAC1* estimated by the ratio of total *HAC1* mRNA to *HAC1*^*i*^ mRNA in *eS7A-WT* and *eS7A-4KR* mutant cells. **h**, Translation efficiency of HAC1, estimated by the ratio of Hac1 protein to *HAC1*^*i*^ mRNA in *eS7A-WT* and *eS7A-4KR* mutant cells. **i**, Translation efficiency after 4 h of Tm treatment, as estimated by ribosome footprints and mRNA reads in the *eS7A-WT* and *eS7A-4KR* mutant cells. **j**, Translation of *HAC1*^*i*^ mRNA is decreased in *eS7A-4KR* mutant cells. Polysome profiles were generated by continuous measurement of absorbance at 254 nm. Equal volumes of the fractions were collected and processed for northern blotting. **k**, Quantification of *HAC1*^*i*^ and *HAC1*^*u*^ mRNA levels in sucrose gradient fractions. **c, f, j**, Cropped blots were display. Cropped blots were display. All uncropped images are available in Supplemental Figure. S8.

**Fig. 3.**
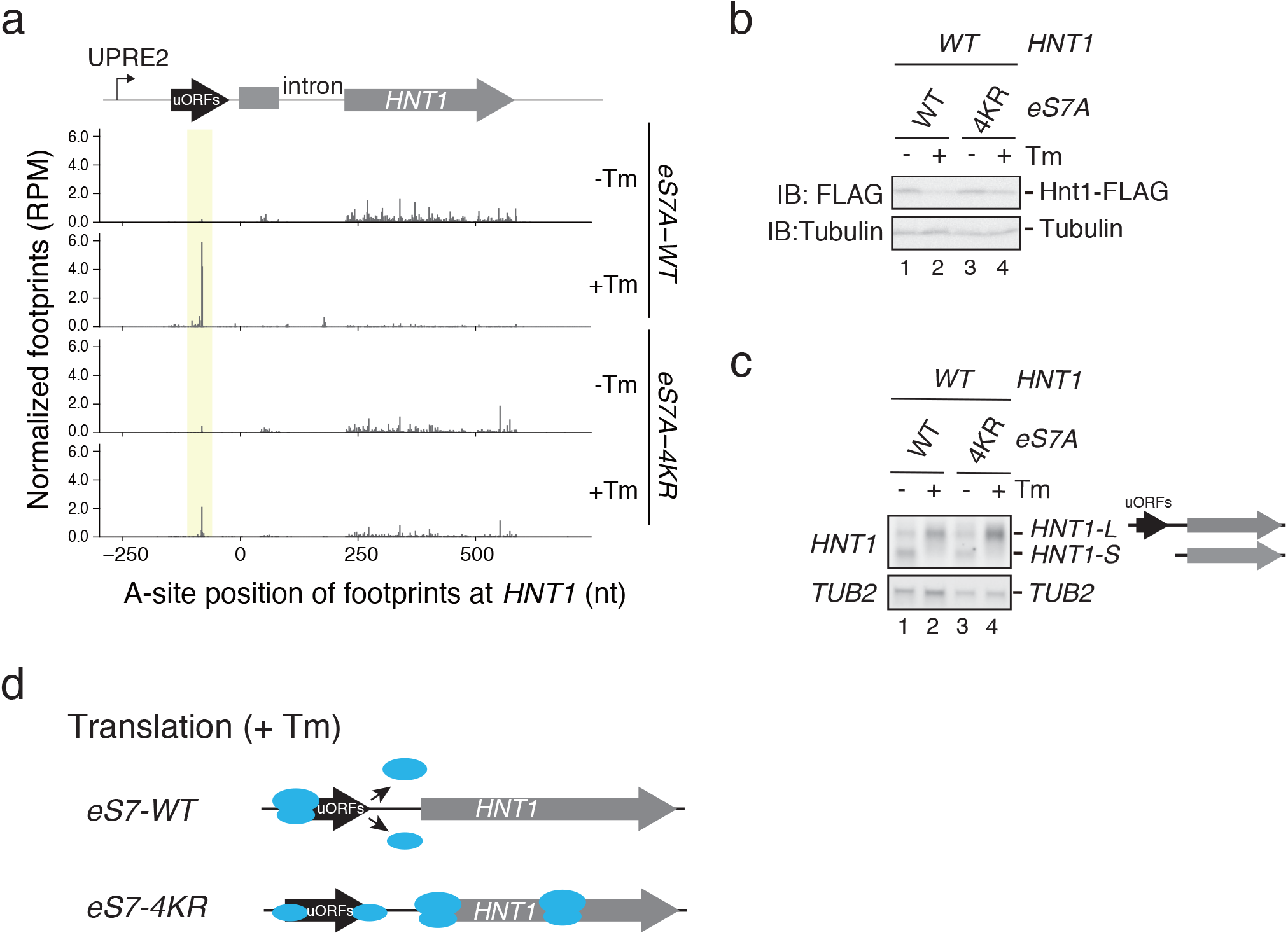
Downregulation of *HNT1* translation during UPR was defective in eS7A-4KR mutant cells. **a**, Map of the A-site position of footprints at *HNT1* in *eS7A-WT* or *eS7A-4KR* mutant cells in the absence (-Tm) or presence of (+Tm) for 4 hr. **b**, Hnt1-FLAG protein level was significantly reduced by Tm treatment. **c**, The uORF-containing *HNT1* mRNA was induced by Tm treatment in *eS7A-WT* and *eS7A-4KR* mutant cells. **d**, Model for uORF-dependent translational downregulation of *HNT1* and the role of eS7 ubiquitination in this regulation. In eS7-WT cells, translation initiation from uORF3 reduced initiation from *HTN1* main ORF in the presence of Tm. In eS7-4KR cells, translation initiation from uORF3 was reduced, and initiation from *HTN1* main ORF was not downregulated upon UOR. **b, c**, Cropped blots were display. All uncropped images are available in Supplemental Figure. S9.

Not4-mediated monoubiquitination of eS7A plays a crucial role in upregulating translation during the UPR, with the translation of *HAC1* most drastically upregulated in response to the UPR. The downregulation of translation of specific mRNAs during the UPR was also abrogated in *eS7A-4KR* mutant cells, with the translation of mRNA encoding Histidine triad NucleoTide-binding 1 (*HNT1)* being most drastically repressed (Fig. 1e). Hac1 induces the expression of long un-decoded transcript isoforms, and downregulation of *HNT1* translation depends on the uORFs, leading to protein downregulation in response to the UPR^37^. Ribosome profiling confirmed the repression of *HNT1* translation, with the TE being downregulated approximately 60-fold 4 h after Tm treatment. Importantly, the UPR-associated downregulation of *HNT1* was significantly diminished in *eS7A-4KR* mutant cells (4-fold; Supplementary Fig. 2d), despite its level of mRNA being slightly increased (Supplementary Fig. 2d).

### Not4-mediated eS7A monoubiquitination is required for translational regulation of HAC1^i^ and HNT1 mRNA

We identified *Hac1* as the most significantly upregulated gene during the UPR (Fig. 1d-e). *HAC1*^*u*^ mRNA is stored in the cytoplasm in the absence of ER stress, and its translation is tightly suppressed by a base-pairing interaction between the intron and the 5′untranslated region (5′UTR)^13,14,38^. Excision of the intron by Ire1-dependent splicing in response to ER stress leads to robust translation of *HAC1*^*i*^ mRNA, with the resulting Hac1 protein upregulating UPR target gene expression. Tm drastically and rapidly increased the TE of *HAC1*^*i*^ mRNA (Fig. 2a; Supplementary Fig. 12a). This increase was less robust in *eS7A-4KR* mutant than in *eS7A-WT* cells (93.05-fold vs. 5.89-fold, respectively; Fig. 2a). Mapping of the footprints throughout *HAC1*^*i*^ mRNA in *eS7A-WT* and *eS7A-4KR* mutant cells (Fig. 2b) suggested that this mRNA does not contain a strong translation pause site.

To validate these results by ribosome profiling, the expression of *HAC1* was evaluated at the splicing, translation and protein levels. Hac1 protein was detected 0.5 h after the addition of Tm to WT cells, but its signal was much weaker in *not4*Δ and *eS7A-4KR* mutant cells, being approximately 50% of the levels in *eS7A-WT* cells (Fig. 2c). The level of Hac1 protein normalized to that of tubulin also indicated that Hac1 protein was significantly downregulated in the *eS7A-4KR* mutant, being approximately 50% of the levels in *eS7A-WT* cells 4 h after Tm treatment (Fig. 2d). In *eS7A-4KR* mutant cells, induction of *HAC1* mRNA by Tm treatment was intact (Fig. 2e-g), although this induction was moderately delayed by about 30 min after the addition of Tm (Fig. 2f-g). The splicing efficiency (the ratio of *HAC1*^*u*^ + *HAC1*^*i*^ mRNAs to *HAC1*^*i*^ mRNA) was moderately reduced in *not4*Δ and *eS7A-4KR* mutant cells (Fig. 2g).

In comparing TE in eS7-WT and eS7-4KR cells 4 h after the addition of Tm based on reads of *HAC1*, we found that the translation of *HAC1* mRNAs was approximately 4-fold lower in eS7-4KR than in wild-type (WT) cells (Fig. 2i). To validate TE by other methods, we analyzed the distribution of *HAC1* mRNA by sucrose gradient ultracentrifugation (Fig. 2j-k). In the absence of Tm, the *HAC1*^*u*^ mRNAs were detected on polysomes, with their distribution and levels being similar in eS7A-4KR and eS7A-WT cells (Fig. 2f, j, k). In the absence of Tm treatment, however, almost no footprints of ribosomes on *HAC1*^*u*^ were observed in eS7A-4KR and eS7A-WT cells (Fig. 2b), consistent with previous findings. These results may have been due to the tightly packed configuration of ribosomes on *HAC1*^*u*^ mRNA. After Tm treatment for 4 h, the levels of *HAC1*^*i*^ mRNA on polysomes were moderately but significantly lower in eS7A-4KR than in eS7A-WT cells (Fig. 2b, k). Overall, the changes in the levels of *HAC1*^*i*^ mRNA on polysomes were consistent with TE calculated based on the levels of *HAC1*^*i*^ mRNA and Hac1 protein, or approximately 2-fold (Fig. 2h). The imperfect correlation of TE with the ratio of protein to mRNA level may be due to a difference in the efficiency of recovery of mRNA reads from ribosomes translating *HAC1*^*i*^ mRNA in eS7A-WT and eS7A-4KR mutant cells.

Initiation of the translation of *HAC1*^*u*^ mRNA has been reported to be tightly suppressed by base-pairing interactions between its intron and 5’UTR, resulting in the stalling of ribosomes in a tightly packed configuration. This may prevent nuclease cleavage during library preparation, resulting in the loss of ribosome footprints derived from *HAC1*^*u*^ mRNA^39^. Therefore, TE calculated from footprints may imperfectly reflect the density of elongating ribosomes on *HAC1*^*u*^ mRNAs. We estimated the TE of *HAC1*^*u*^ mRNAs based on their distribution in the sucrose density gradients (Fig. 2j-k). In the absence of Tm, *HAC1*^*u*^ mRNAs were detected on the polysomes, and their distribution was similar in eS7A-4KR and eS7A-WT cells (Fig. 2j, k, top panel). Although the levels of *HAC1*^*u*^ mRNA were almost the same in the eS7A-4KR and eS7A-WT cells (Fig. 2f), almost no footprints from ribosomes on *HAC1*^*u*^ were observed in eS7A-WT and eS7A-4KR cells in the absence of Tm treatment (Fig. 2b), suggesting that ribosomes form tightly packed configuration on *HAC1*^*u*^ mRNA in S7A-WT and eS7A-4KR cells.

Not4 monoubiquitinates eS7 at four lysine residues (Supplementary Fig. 4a), with K83 ubiquitination being primarily responsible for mRNA quality control^24^. To identify the ubiquitination site(s) of eS7A required for the upregulation of *HAC1*^*i*^, *HAC1*^*u*^ mRNA splicing and Hac1 protein levels were examined in four mutants containing a single lysine residue, eS7A, susceptible to Not4-mediated monoubiquitination, *eS7A-3KR-K72, eS7A-3KR-K76, eS7A-3KR-K83* and *eS7A-3KR-K84*^24^. Hac1 protein levels were significantly lower in *eS7A-3KR-K72* and *eS7A-3KR-K76* single-lysine and *eS7A-4KR* mutant cells (Supplementary Fig. 4b, d), but not in *eS7A-3KR-K83* and *eS7A-3KR-K84* single-lysine cells (Supplementary Fig. 4c, e). These results, which indicate that monoubiquitination of eS7A at lysine 83 or 84 is sufficient for the production of Hac1 protein, are consistent with a model in which translation of *HAC1*^*i*^ mRNA is facilitated by monoubiquitinated eS7A at lysine residue 83 or 84. However, the possibility that mutation of eS7A could cause structural changes to the ribosome in addition to the ubiquitination defect cannot be excluded.

**Fig. 4.**
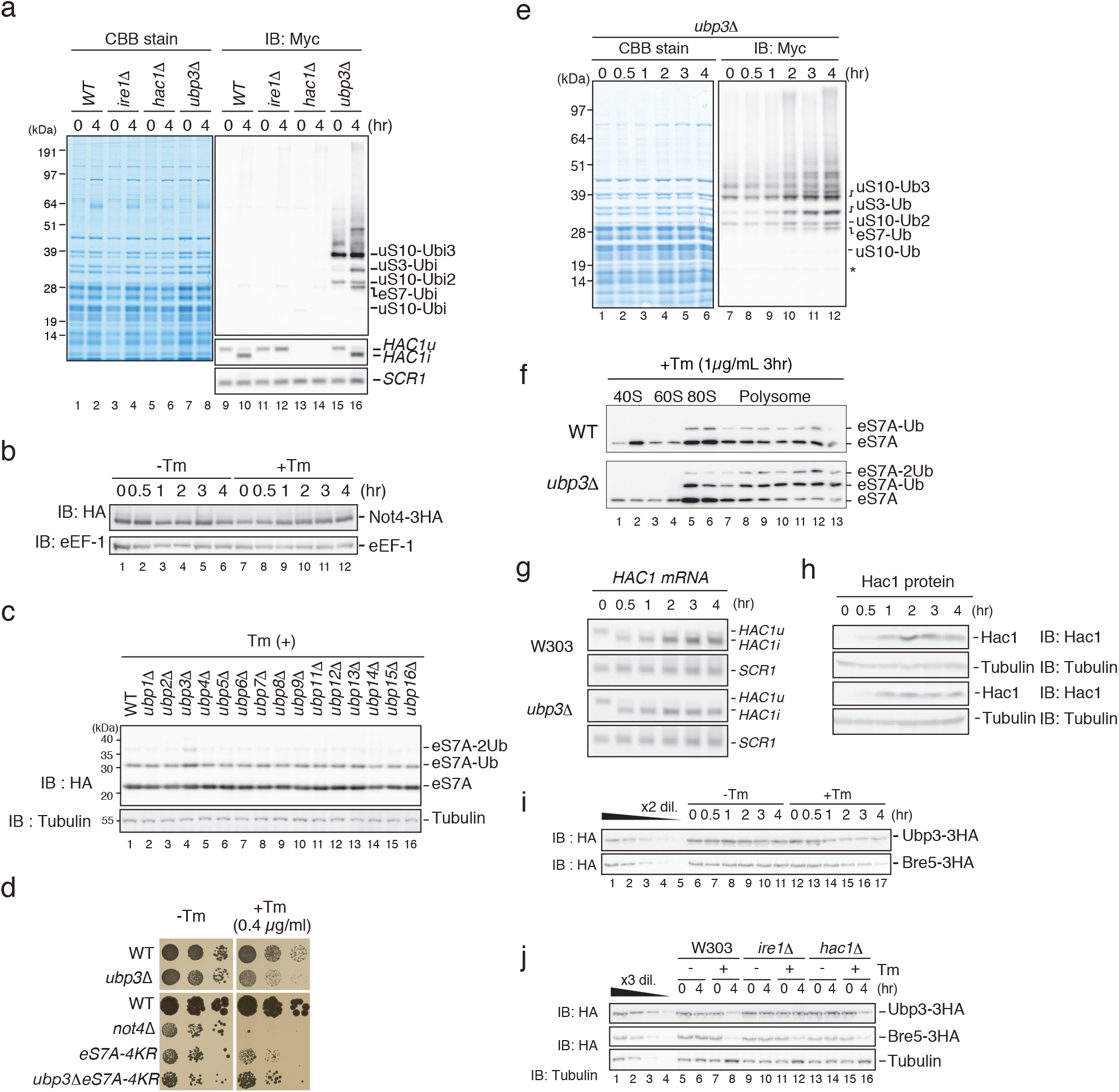
Deubiquitinating enzyme complex Upb3-Bre5 is involved in the regulation of eS7A ubiquitination during UPR. **a**, Regulation of eS7A ubiquitination during UPR was independent of Ire1 and Hac1. **b**, UPR had no effect on Not4 levels. **c**, A genetic screen to identify the enzyme response for the deubiquitination of polyubiquitinated S7A. The levels of polyubiquitinated eS7A 2 h after Tm addition were significantly and specifically increased in *ubp3*Δ mutant cells. **d**, Ubp3 is a deubiquitinating enzyme of eS7A and is required for Tm resistance. **e**, Levels of polyubiquitinated eS7A during UPR in *ubp3*Δ mutant cells. **f**, Monosomes and polysomes, but not free 40S, in both wild-type and *ubp3*Δ mutant cells contain polyubiquitinated eS7A. **g**, The splicing of *HAC1*^*u*^ mRNA was intact in *ubp3*Δ mutant cells. **h**, Hac1 protein level was slightly reduced in *ubp3*Δ mutant cells. **i**, Ubp3 and its co-factor Bre5 were significantly and gradually decreased as a function of time during UPR. **j**, Downregulation of Ubp3 but not Bre5 was impaired in *ire1*Δ cells, but not in *hac1*Δ cells. **a, b, c, e, f, j, h, i, j**, Cropped gels or blots were display. All uncropped images are available in Supplemental Figure. S10.

After Tm treatment, the translation of *HNT1* mRNA is suppressed in an uORF-dependent manner^37,40^. We also found that *HNT1* was most markedly repressed upon UPR (Fig. 1e). Hac1 is reported to be required to synthesize uORF-containing *HNT1* mRNAs, making it essential for the downregulation of *HNT1*. Interestingly, assessment of ribosome occupancy on *HNT1* mRNA (Fig. 3a) showed that the ribosomes efficiently read through the uORF in the eS7A-4KR mutant after Tm treatment. The reduction of Hnt1 protein after Tm treatment was significantly restored in the eS7A-4KR mutant (Fig. 3b), but the level of long transcripts containing uORFs was not reduced in the mutant (Fig. 3c). These findings indicate that, although the level of Hac1 was lower in eS7A-4KR mutant than in es7A-WT cells, it was sufficient in the former to induce uORF-containing *HNT1* mRNA under UPR conditions. Thus, independently of Hac1, eS7 ubiquitination may facilitate the translation of uORFs, thereby repressing the translation of *HNT1* ORF (Fig. 3d).

### Deubiquitinating enzyme complex Upb3-Bre5 is involved in the regulation of eS7A ubiquitination during UPR

Our results demonstrated that eS7A ubiquitination is required for translational controls during UPR. We therefore assessed whether the upregulation of eS7A ubiquitination is dependent on Ire1 and Hac1. We found that eS7A ubiquitination was increased in *ire1*Δ and *hac1*Δ mutant cells 4 hr after Tm addition (Fig. 4a), indicating that the Ire1-Hac1 pathway is not required for the upregulation of eS7A ubiquitination.

To further elucidate the mechanism underlying the regulation of eS7A ubiquitination, we measured the expression of the E3 ligase Not4, which is responsible for the monoubiquitination of eS7A. We found that Not4 expression remained unchanged after the addition of Tm (Fig. 4b). We next hypothesized that the increase in monoubiquitinated eS7A in response to Tm is caused by a decrease in deubiquitinating activity. To test this possibility, we first performed a genetic screen to identify the enzyme responsible for deubiquitinating ubiquitinated eS7A. The levels of ubiquitinated eS7A were significantly and specifically increased in *ubp3*Δ mutant cells 2 h after the addition of Tm (Fig. 4c, lane 4). Deletion of *UBP3* conferred sensitivity to Tm (Fig. 4d), suggesting that Ubp3 is the enzyme responsible for deubiquitinating ubiquitinated eS7A and that it contributes to resistance to ER stress. To measure the levels of ubiquitinated eS7A during UPR, we used N-terminal Myc-tagged ubiquitin, followed by ribosome affinity purification^22^. Tm addition significantly upregulated monoubiquitinated eS7A in a time-dependent manner (Fig. 4e). Western blot analysis of sucrose gradient fractions showed that monosomes or polysomes, but not free 40S subunits, contained ubiquitinated eS7A in both WT and *ubp3*Δ mutant cells (Fig. 4f), suggesting that translating ribosomes contain polyubiquitinated eS7A. In *ubp3*Δ mutant cells, the levels of mono- and di-ubiquitinated eS7A were increased in mono- and polysome fractions, but not in the 40S subunit (Fig. 4f), indicating that eS7A is the substrate of Ubp3 in monosomes and polysomes but not in the 40S subunit. Assessments of the levels of expression of *HAC1* mRNA and Hac1 protein in the *ubp3*Δ deletion mutant showed that Hac1 protein expression was significantly decreased (Fig. 4g-h).

Next, we assessed whether the UPR results in the downregulation of the deubiquitinating enzyme. Ubp3 forms a complex with its cofactor Bre5 *in vivo*, with complex formation required for Ubp3 function^41,42^. Western blot analysis of Ubp3-3HA and Bre5-3HA expressed from endogenous promoters showed that the levels of the deubiquitinating enzyme Ubp3 and its cofactor Bre5 decreased significantly and gradually during the UPR (Fig. 4i). The downregulation of Ubp3 was impaired in *ire1*Δ but not in *hac1*Δ mutant cells, although neither of these UPR factors was required for the downregulation of Bre5 (Fig. 4j). These results indicate that activated Ire1 induces the downregulation of Ubp3, but not of Bre5, during the UPR. Taken together, these findings indicate that downregulation of deubiquitination in response to UPR increases the levels of monoubiquitinated eS7A, a downregulation that is affected by, but not completely dependent on, the Ire1-Hac1 pathway (Fig. 5).

**Fig. 5.**
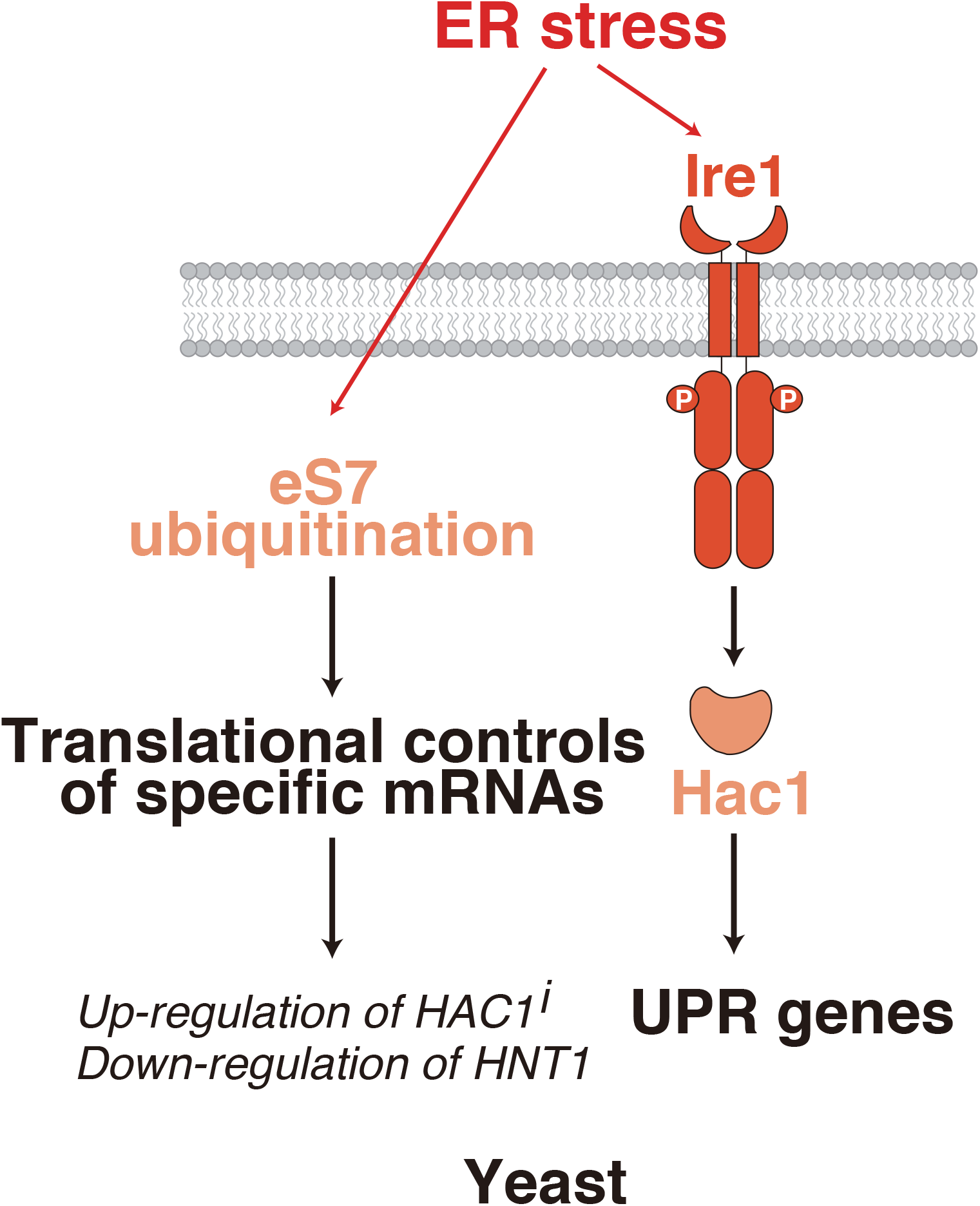
Model for regulation of eS7A ubiquitination in response to UPR and its roles in translational controls. Not4-mediated monoubiquitination of eS7A at lysine 83 or 84 is required to control translation during UPR. Tm-induced ER stress increased the levels of ubiquitinated eS7A in a manner independent of Ire1 and Hac1. Monoubiquitinated eS7A is required for upregulation of specific mRNAs including *HAC1*^*i*^ mRNA and downregulation of *HNT1* mRNA. Ribosome ubiquitination of eS7A is therefore likely required for translational control in response to ER stress in yeast.

## DISCUSSION

Ubiquitination of ribosomal proteins plays an essential role in quality control induced by ribosomal stalling^22-24,26,28,37^. However, the physiological functions of ribosome modifications remain unclear. The findings of this study indicate that ribosome eS7 monoubiquitination is required for translational controls during ER stress responses in yeast. Ribosome profiling revealed that eS7A monoubiquitination was necessary for translational up- and down-regulation of specific mRNAs. Monoubiquitination of eS7A facilitated the translational upregulation of *HAC1*^*i*^ mRNA, a master transcription factor in the UPR, and the downregulation of *HNT1* during ER stress. The mechanisms underlying the roles of eS7A ubiquitination in translational regulation of specific mRNAs during ER stress, however, remain unclear. Hac1-mediated production of long transcripts containing uORFs was shown to repress the translation of histidine triad nucleotide-binding 1 (*HNT1*) mRNA^37^. We recently reported that uORF3 is required for *HNT1* expression, and that translation of *HNT1* is efficiently repressed by a strong Kozak sequence uORF3 during UPR^43^. These findings suggest that initiation of translation at the AUG codon of uORF3 is inefficient, and that leaky scanning of uORF3 is responsible for translation of *HNT1*. Although Tm treatment reduced the production of Hnt1 protein in eS7A-WT, it did not alter Hnt1 protein production in the eS7A-4KR mutant (Fig. 3b), despite the level of long transcripts containing uORFs not being changed in this mutant (Fig. 3c). These findings suggested that initiation of translation at the AUG codon of uORF3 is repressed in the eS7A-4KR mutant, and that initiation of translation at the AUG codon of the *HNT1* ORF is stimulated by the increase in leaky scanning of uORF3. Thus, eS7 ubiquitination may facilitate the translation of uORF3, thereby repressing the translation of *HNT1* ORF.

Understanding translational regulation in response to the accumulation of unfolded proteins in the ER can be improved by determining the molecular mechanisms underlying eS7A monoubiquitination-mediated regulation of translation during the UPR. Our results suggested that initiation of translation of specific mRNAs, including *HAC1*^*i*^ and uORF3 of *HNT1* mRNA, depends on eS7 ubiquitination by an as yet unknown mechanism, and that reduced translation from these initiation codons resulted in defects in the upregulation of *HAC1*^i^ mRNA and the downregulation of *HNT1* mRNA upon UPR (Fig. 5). These interactions between eS7A and translation initiation factors may be critical for initiating translation at specific sites. eIF3 binding to ribosomes elongating and terminating on short upstream ORFs has been shown to promote the re-initiation of *GCN4* translation^44-47^. Moreover, eIF3-dependent translation initiation mechanism contributes to translational recovery in chronic ER stress response^7^.The recently resolved cryo-EM structure of eIF3 in the context of the human 43S pre-initiation complex^46,48,49^, and the proximity of eIF3 to eS7A in a yeast 48S pre-initiation complex model suggest that eS7A is associated with specific initiation factors^49^ (Supplementary Fig. 5). Modification of eS7A, including ubiquitination, may affect the interaction of the 40S subunit with translation initiation factors in the pre-initiation complex, modulating the initiation of translation of specific mRNAs.

The level of monoubiquitinated eS7A in response to UPR was also upregulated in *ire1*Δ and *hac1*Δ mutant cells, indicating that the Ire-Hac1 pathway is not necessary for the regulation of eS7A ubiquitination. These results strongly suggest that the deubiquitinating enzyme complex Upb3-Bre5 was involved in the regulation of eS7A ubiquitination. The level of monoubiquitinated eS7A was upregulated in *ubp3*Δ mutant cells. Ubp3 is the enzyme that deubiquitinates ubiquitinated eS7A and contributes to cell resistance to ER stress. Bre5 is a regulatory subunit that is downregulated upon UPR, even in *ire1*Δ and *hac1*Δ mutant cells, indicating that the downregulation of the deubiquitinating enzyme complex Ubp3-Bre5 is independent of the Ire1-Hac1 pathway. These findings suggest that ER stress reduced the levels of the deubiquitinating enzyme complex Ubp3-Bre5, leading to an increase in the ubiquitinated form of eS7A. Monoubiquitinated eS7A facilitates the translation of *HAC1*^*i*^ mRNA, resulting in efficient induction of Hac1-target genes and downregulation of *HNT1* by as yet unknown mechanisms. Further investigations are needed to determine the mechanism underlying the ribosome ubiquitination-mediated regulation of translation initiation upon UPR.

## MATERIALS AND METHODS

### Materials and Methods

#### Yeast strains and genetic methods

The *S. cerevisiae* strains used in this study are listed in Table 1. Gene disruption and C-terminal tagging were performed as previously described^50,51^. *S. cerevisiae* W303-1a based strains were obtained by established recombination techniques using PCR-amplified cassette sequences (*kanMX4, hphMX4, natMX4, natNT2* or *HISMX6*) (Longtine, M. S. *et al*., Janke, C. *et al*.,). To construct strains of essential ribosomal protein genes (uS10, uS3, eS7AeS7B), the shuffle strain transformed with plasmid expressing mutant ribosomal protein products was grown on SDC plates containing 0.5 mg/mL 5-fluoroorotic acid (5-FOA, #F9001-5, Zymo Research) and a strain lacking URA3 was isolated.

#### Plasmid constructs

Specific DNA sequences were PCR amplified using gene specific primers and cloned into vectors using PrimeSTAR HS DNA polymerase (#R010A, Takara-bio) and T4 DNA ligase (#M0202S, NEB). The sequences of all cloned DNAs amplified by PCR were verified by sequencing. Plasmids and primers used in this study are listed in Tables 2 and 3, respectively.

#### Yeast Culture and Media

All yeast cells were cultured at 30°C in YPD or synthetic complete (SC) medium containing 2% glucose and harvested during log phase by centrifugation. To induce ER stress, yeast cells were grown at 30°C until their absorbance at 600 nm was 0.2, treated with 1 µg/mL tunicamycin (Tm, #208-08243, Wako) for ∼4 h and harvested. For polysome analysis, yeast cells cultured at 30°C for 3 h after Tm addition were treated for 5 min with 0.1 mg/mL cycloheximide (CHX, #06741-04, Nacalai tesque) before harvesting by centrifugation. All cell pellets were frozen in liquid nitrogen immediately after harvest and stored at -80°C until used.

#### RNA Isolation

Total RNA was isolated from yeast cells by acidic phenol RNA extraction (Ikeuchi and Tesina et al., 2019) with several modifications. Each cell pellet was re-suspended on ice in 200 µL RNA buffer (300 mM NaCl, 20 mM Tris-HCl (pH 7.5), 10 mM EDTA, 1% SDS, in DEPC-treated MilliQ water), followed by immediate addition of 250 µL of water-saturated phenol. The preparations were mixed well by vortexing for 10 sec, incubated at 65°C for 5 min, again mixed by vortexing for 10 sec and chilled on ice for 5 min. After centrifugation at 16,000 × *g* for 5 min at room temperature, 190 µL of each water layer was transferred to a new 1.5 mL RNase free tube. A 250 µL aliquot of water-saturated phenol was added and the procedure was repeated. After centrifugation, 170 µL of each water layer was transferred to a new 1.5 mL RNase free tube; 200 µL of water-saturated phenol/chloroform (1:1) was added; and the tubes were vortexed for 10 sec and centrifuged at 16,000 × *g* for 5 min at room temperature. A 150 µL aliquot of each water layer was transferred to a new 1.5 mL RNase free tube, to which was added 200 µL of water-saturated phenol/chloroform/isoamylalchol (25:24:1), followed by vortexing for 10 sec and centrifugation at 16,000 × *g* for 5 min at room temperature. Finally, 130 µL of each water layer was transferred to a new 1.5 mL RNase free tube and subjected to ethanol precipitation. Each RNA pellet was dissolved in 20–30 µL of DEPC-treated water.

#### RNA Electrophoresis and Northern Blotting

RNA electrophoresis and northern blotting were performed as described (Ikeuchi and Tesina et al., 2019) with the following modifications. A 6 µL aliquot (1 µg) of total RNA solution was mixed with 24 µL of glyoxal solution [600 µL DMSO, 200 µL deionized 40% glyoxal, 120 µL 10x MOPS buffer (200 mM MOPS, 50 mM NaOAc, 10 mM EDTA, pH 7.0), 62.5 µL of 80% glycerol, and 17.5 µL of DEPC-treated water in 1 μL] and 3 µL of RNA loading buffer (50% glycerol, 10 mM EDTA pH 8.0, 0.05% bromophenol blue and 0.05% xylene cyanol). The mixture was incubated at 74°C for 10 min and left to stand on ice for 10 min. A 25 µL aliquot of each sample was electrophoresed at 200 V for 40 min on a 1.2% agarose gel in 1× MOPS buffer (20 mM MOPS, 5 mM NaOAc and 1 mM EDTA, pH 7.0), followed by capillary transfer of RNA to Hybond-N+ membranes (GE Healthcare) with 20x SSC (3 M NaCl and 300 mM trisodium citrate dihydrate) for 18 h. RNA on the membranes was cross-linked with CL-1000 ultraviolet crosslinker (UVP) at 120 mJ/cm^2^. The membranes were incubated with DIG Easy Hyb Granules (#11796895001, Roche) for 1 hr in a hybridization oven at 50°C. DIG-labelled probe prepared using PCR DIG Probe Synthesis Kit (#11636090910, Roche) was added and the membranes incubated for over 18 h. The membranes were washed twice with wash buffer I (2x SSC, 0.1% SDS) for 15 min each in a hybridization oven at 50°C, washed once with wash buffer II (0.1x SSC, 0.1% SDS) for 15 min at 50°C, and washed once with 1x maleic acid buffer (100 mM maleic acid, 150 mM NaCl, pH 7.0, adjusted by NaOH) for 10 min at room temperature. The membranes were incubated with Blocking Reagent (#11096176001, Roche) for 30 min and then with anti-Digoxigenin-AP, Fab fragments (#11093274910, Roche) in Blocking Reagent for 1 h. The membranes were washed three times with wash buffer III (1x maleic acid buffer, 0.3% tween 20) for 10 min each, incubated in equilibration buffer (100 mM Tris-HCl, 100 mM NaCl, pH 9.5), and reacted with CDP-star (#11759051001, Roche) for 10 min. Chemiluminescence was detected using LAS-4000 (GE Healthcare), and relative RNA levels were determined using Multi Gauge v3.0 (Fujifilm, Japan).

#### Trichloroacetic acid (TCA) precipitation for protein preparation

Yeast cell pellets in 1.5 mL tubes were resuspended in 500 µL ice-cold TCA buffer (20 mM Tris-HCl pH 8.0, 50 mM NH_4_OAc, 2 mM EDTA, and 1 mM PMSF) and transferred to new 1.5 mL tubes containing 500 µL 20% TCA and 500 µL 0.5 mm Zirconia/Silica Beads (BioSpec). The mixtures were vortexed three times for 30 sec each, and the supernatants were transferred to new 1.5 mL tubes. A 500 µL aliquot of ice-cold TCA buffer was added to each tube, followed by vortexing for 30 sec and transfer of the supernatant to a new 1.5 mL tube. The lysates were centrifuged at 14,000 rpm for 15 min at 4°C), the supernatants were discarded, and each pellet was resuspended in SDS sample buffer (125 mM Tris-HCl pH 6.8, 4% SDS, 20% glycerol, 100 mM DTT, and 0.01% bromophenol blue) and heated at 95°C for 5 min or at 65°C for 15 min. These samples were subsequently loaded onto SDS-PAGE or Nu-PAGE gels.

#### Ribosome purification to observe ribosome ubiquitination

To assess ribosome ubiquitination during ER stress, ribosomes were purified with Myc-tagged ubiquitin (Myc-Ubi) and the FLAG-tagged ribosomal protein uL23 (uL23-FLAG), as described previously (Matsuo and Ikeuchi et al., 2017). Yeast cells harbouring p*CUP1*p-*MYC-UBI* and p*RPL25(uL23)-FLAG* were cultured in 800 mL of synthetic complete medium. To induce the expression of Myc-Ubi, the cells were cultured in the presence of 0.1 mM Cu^2+^ for 1 h, followed by the addition of Tm to a concentration of 1 µg/mL and harvesting at the indicated time points. Cell lysates were prepared, and FLAG-tagged ribosomes were purified using M2 FLAG-affinity resin (Sigma), as described previously (Inada et al., 2002). Affinity purified samples were subjected to SDS-PAGE followed by staining with Coomassie brilliant blue (CBB) or western blotting with an anti-Myc antibody.

#### Protein Electrophoresis and Western Blotting

Protein electrophoresis and western blotting were performed as described previously (Ikeuchi and Tesina et al., 2019) with the following modifications. Protein samples were separated by SDS-PAGE or Nu-PAGE, and stained with CBB or transferred to PVDF membranes (Immobilon-P, Millipore). After blocking with 5% skim milk in PBST (10 mM Na_2_HPO_4_/NaH_2_PO_4_ pH 7.5, 0.9% NaCl, and 0.1% Tween-20), the membranes were incubated with primary antibodies (Table 4) for 1 h at room temperature followed by three washes with PBST and further incubation with horseradish peroxidase (HRP)-conjugated secondary antibodies for 1 h at room temperature. If detecting HA-tagged proteins, the membranes were incubated with HRP-conjugated antibodies. After three washes with PBST, chemiluminescence was detected by LAS4000 (GE Healthcare).

#### Sucrose density gradient (SDG) centrifugation

SDG was performed as described (Matsuo and Ikeuchi, 2017) with the following modifications. Yeast cells were grown exponentially at 30°C and treated with 0.1 mg/mL cycloheximide for 5 min before harvesting by centrifugation. The cell pellets were frozen and ground in liquid nitrogen using a mortar and pestle. The cell powder was resuspended in lysis buffer (20 mM HEPES-KOH, pH 7.4, 100 mM potassium acetate, 2 mM magnesium acetate, 0.5 mM dithiothreitol, 1 mM phenylmethylsulfonyl fluoride; Complete mini EDTA-free; #11836170001, Roche) to prepare the crude extracts. Sucrose gradients (10–50% sucrose in 10 mM Tris-acetate, pH 7.4, 70 mM ammonium acetate, and 4 mM magnesium acetate) were prepared in 25 × 89 mm polyallomer tubes (Beckman Coulter) using a Gradient Master (BioComp). Crude extracts (the equivalent of 50 A260 units) were layered on top of the sucrose gradients, followed by centrifugation at 150,000 ×g in a P28S rotor (Hitachi Koki, Japan) for 2.5 h at 4°C. The gradients were fractionated (BioComp Piston Gradient Fractionator), and the polysome profiles generated by continuous measurement of absorbance at 254 nm using a single path UV-1 optical unit (ATTO Biomini UV-monitor) connected to a chart recorder (ATTO digital mini-recorder). For western blotting of these fractions, 900 µL of each fraction were mixed with 180 µL of 100% TCA and the mixtures incubated for 15 min at 4°C. After centrifugation at 14,000 rpm for 15 min at 4°C, the supernatants were removed, and each pellet was dissolved in SDS sample buffer (125 mM Tris-HCl pH 6.8, 4% SDS, 20% glycerol, 100 mM DTT, and 0.01% bromophenol blue) and heated at 65°C for 15 min.

#### Spot assay

Yeast cells were cultured in 2 mL YPD or SC medium containing 2% glucose at 30°C for 12– 24 hr and adjusted to an optical density at 600 nm of 0.3. Ten-fold serial dilutions were prepared in 1.5 mL tubes and spotted onto plates with and without Tm. The plates were incubated at 30°C for 2–3 days.

#### Ribosome profiling and RNA-seq

To induce ER stress, yeast cells were grown at 30°C until reaching an optical density at 600 nm of 0.2. The cells were treated with 1 µg/mL Tm for ∼4 h, harvested by vacuum filtration, and lysed by cryogenic grinding in a mixer mill (Retsch MM400). Whole cell lysates containing 10 µg of total RNA were each treated with 12.5 units of RNase I (Epicentre) at 23°C for 45 min, and the ribosome fraction was sedimented through a 1 M sucrose cushion. The ribosome protected mRNA fragments were extracted with TRIzole regent (Life Technologies) and used for library preparation.

Library preparation was performed as described (McGlincy and Ingolia, 2017) with the following modifications. For ribosome profiling analysis, whole cell lysates containing 20 µg of total RNA were each treated with 10 units of RNase I (Epicentre) at 24°C for 45 min. Linker DNA consisted of 5’-(Phos)NNNNNIIIIITGATCGGAAGAGCACACGTCTGAA(ddC)-3’, with (Phos) indicating 5’ phosphorylation; (ddC) indicating a terminal 2’, 3’-dideoxycytidine; and Ns and Is indicate random barcodes for eliminating PCR duplication and multiplexing barcodes, respectively. The linkers were pre-adenylated with a 5’ DNA Adenylation kit (NEB), and then used for the ligation reaction. Un-reacted linkers were digested with 5’ deadenylase (NEB) and RecJ exonuclease (epicentre) at 30°C for 45 min. RNA was reverse transcribed using the oligonucleotide primer, 5’-(Phos)NNAGATCGGAAGAGCGTCGTGTAGGGAAAGAG (iSp18)GTGACTGGAGTTCAGACGTGTGCTC-3’. PCR was performed with the primers, 5’-AATGATACGGCGACCACCGAGATCTACACTCTTTCCCTACACGACGCTC-3’ and 5’-CAAGCAGAAGACGGCATACGAGATJJJJJJGTGACTGGAGTTCAGACGTGTG-3’, where Js indicate reverse complement of the index sequence determined during Illumina sequencing. For RNA-seq analysis, total RNA was extracted from lysate using Trizol reagent (Life Technologies); rRNAs were removed from the total RNA using the Ribo-Zero Gold rRNA Removal Kit (Yeast) (Illumina); and the cDNA libraries were prepared using a TruSeq Stranded mRNA Library Prep Kit (Illumina). The libraries were sequenced on a HiSeq 2000/4000 (Illumina). The reads were mapped to yeast transcriptome, removing duplicated reads based on random barcode sequences. The analyses for ribosome profiling were restricted to read lengths of 30–33 nt for eS7-WT (0 h after Tm treatment), 29–33 nt for eS7-WT (4 h after Tm treatment), and 28–32 nt for eS7A-4KR datasets. The position of the A-site from the 5’-end of the reads was estimated based on the length of each footprint. The offsets using for analysis of ribosome profiling were 17 for 32–33 nt, 16 for 29–31 nt and 15 for 28 nt reads. For analysis of RNA-seq, an offset 15 was used for all mRNA fragments. The DESeq package was used to calculate the fold change of mRNA expression and TE (Anders et al 2010).

## Acknowledgments

The authors thank the members of their laboratories for discussions and critical comments on the manuscript. This study was supported by a Grant-in-Aid for Scientific Research (KAKENHI) from the Japan Society for the Promotion of Science (Grant numbers 18H03977 and 19H05281 to T.I., and 19K06481 to Y.M.), and the Uehara Memorial Foundation (to T.I.) and the Suzuken Memorial Foundation (to T.U.).

## Author Contributions

Genetic and biochemical experiments were performed by Ya.M., Y.N., and H.Y., under the supervision of T.I. S.I, Yo.M., and T.U. Yo.M. performed the ribosome profiling experiments, under the supervision of N.I. and T.I. Y.S. performed mass spectrometry under the supervision of K.T., T.I. Ya.M., Yo.M., and T.Y., S.I., S.Li. and T.I. wrote the manuscript. T.I. primarily conceived the idea and designed the experiments.

## References

1 Gardner, B. M., Pincus, D., Gotthardt, K., Gallagher, C. M. & Walter, P. Endoplasmic reticulum stress sensing in the unfolded protein response. Cold Spring Harb Perspect Biol 5, a013169, doi: 10.1101/cshperspect.a013169 (2013).

2 Hollien, J. et al. Regulated Ire1-dependent decay of messenger RNAs in mammalian cells. J Cell Biol 186, 323–331, doi: 10.1083/jcb.200903014 (2009).

3 Yoshida, H., Matsui, T., Yamamoto, A., Okada, T. & Mori, K. XBP1 mRNA is induced by ATF6 and spliced by IRE1 in response to ER stress to produce a highly active transcription factor. Cell 107, 881–891 (2001).

4 Adachi, Y. et al. ATF6 is a transcription factor specializing in the regulation of quality control proteins in the endoplasmic reticulum. Cell Struct Funct 33, 75–89 (2008).

5 Harding, H. P., Zhang, Y. & Ron, D. Protein translation and folding are coupled by an endoplasmic-reticulum-resident kinase. Nature 397, 271–274, doi: 10.1038/16729 (1999).

6 Harding, H. P. et al. Regulated translation initiation controls stress-induced gene expression in mammalian cells. Mol Cell 6, 1099–1108 (2000).

7 Guan, B. J. et al. A Unique ISR Program Determines Cellular Responses to Chronic Stress. Mol Cell 68, 885-900.e886, doi: 10.1016/j.molcel.2017.11.007 (2017).

8 Cox, J. S., Shamu, C. E. & Walter, P. Transcriptional induction of genes encoding endoplasmic reticulum resident proteins requires a transmembrane protein kinase. Cell 73, 1197–1206 (1993).

9 Mori, K., Ma, W., Gething, M. J. & Sambrook, J. A transmembrane protein with a cdc2+/CDC28-related kinase activity is required for signaling from the ER to the nucleus. Cell 74, 743–756 (1993).

10 Gardner, B. M. & Walter, P. Unfolded proteins are Ire1-activating ligands that directly induce the unfolded protein response. Science 333, 1891–1894, doi: 10.1126/science.1209126 (2011).

11 Pincus, D. et al. BiP binding to the ER-stress sensor Ire1 tunes the homeostatic behavior of the unfolded protein response. PLoS Biol 8, e1000415, doi: 10.1371/journal.pbio.1000415 (2010).

12 Korennykh, A. V. et al. The unfolded protein response signals through high-order assembly of Ire1. Nature 457, 687–693, doi: 10.1038/nature07661 (2009).

13 Aragon, T. et al. Messenger RNA targeting to endoplasmic reticulum stress signalling sites. Nature 457, 736–740, doi: 10.1038/nature07641 (2009).

14 Ruegsegger, U., Leber, J. H. & Walter, P. Block of HAC1 mRNA translation by long-range base pairing is released by cytoplasmic splicing upon induction of the unfolded protein response. Cell 107, 103–114 (2001).

15 Travers, K. J. et al. Functional and genomic analyses reveal an essential coordination between the unfolded protein response and ER-associated degradation. Cell 101, 249–258 (2000).

16 Krishnan, K. et al. Polysome profiling reveals broad translatome remodeling during endoplasmic reticulum (ER) stress in the pathogenic fungus Aspergillus fumigatus. BMC Genomics 15, 159, doi: 10.1186/1471-2164-15-159 (2014).

17 Ito-Harashima, S., Kuroha, K., Tatematsu, T. & Inada, T. Translation of the poly(A) tail plays crucial roles in nonstop mRNA surveillance via translation repression and protein destabilization by proteasome in yeast. Genes Dev 21, 519–524, doi: 10.1101/gad.1490207 (2007).

18 Dimitrova, L. N., Kuroha, K., Tatematsu, T. & Inada, T. Nascent peptide-dependent translation arrest leads to Not4p-mediated protein degradation by the proteasome. J Biol Chem 284, 10343–10352, doi: 10.1074/jbc.M808840200 (2009).

19 Bengtson, M. H. & Joazeiro, C. A. Role of a ribosome-associated E3 ubiquitin ligase in protein quality control. Nature 467, 470–473, doi: 10.1038/nature09371 (2010).

20 Brandman, O. et al. A ribosome-bound quality control complex triggers degradation of nascent peptides and signals translation stress. Cell 151, 1042–1054, doi: 10.1016/j.cell.2012.10.044 (2012).

21 Defenouillere, Q. et al. Cdc48-associated complex bound to 60S particles is required for the clearance of aberrant translation products. Proc Natl Acad Sci U S A 110, 5046–5051, doi: 10.1073/pnas.1221724110 (2013).

22 Matsuo, Y. et al. Ubiquitination of Stalled Ribosome Triggers Ribosome-associated Quality Control. Nature Communications, doi: 10.1038/s41467-017-00188-1 (2017).

23 Sitron, C. S., Park, J. H. & Brandman, O. Asc1, Hel2, and Slh1 couple translation arrest to nascent chain degradation. RNA 23, 798–810, doi: 10.1261/rna.060897.117 (2017).

24 Ikeuchi, K. T., P.; Matsuo, Y.; Sugiyama, T.; Cheng, J.; Becker, T.; Beckmann, R. and Inada, T. Collided ribosomes form a unique structural interface to induce Hel2-driven quality control pathways. EMBO Journal e100276, doi: 10.15252/embj.2018100276 (2019).

25 Juszkiewicz, S. & Hegde, R. S. Initiation of Quality Control during Poly(A) Translation Requires Site-Specific Ribosome Ubiquitination. Mol Cell 65, 743–750 e744, doi: 10.1016/j.molcel.2016.11.039 (2017).

26 Sundaramoorthy, E. et al. ZNF598 and RACK1 Regulate Mammalian Ribosome-Associated Quality Control Function by Mediating Regulatory 40S Ribosomal Ubiquitylation. Mol Cell 65, 751–760 e754, doi: 10.1016/j.molcel.2016.12.026 (2017).

27 Garzia, A. et al. The E3 ubiquitin ligase and RNA-binding protein ZNF598 orchestrates ribosome quality control of premature polyadenylated mRNAs. Nat Commun 8, 16056, doi: 10.1038/ncomms16056 (2017).

28 Juszkiewicz, S. et al. ZNF598 Is a Quality Control Sensor of Collided Ribosomes. Mol Cell 72, 469–481 e467, doi: 10.1016/j.molcel.2018.08.037 (2018).

29 Sugiyama, T. et al. Sequential Ubiquitination of Ribosomal Protein uS3 Triggers the Degradation of Non-functional 18S rRNA. Cell Rep 26, 3400-3415.e3407, doi: 10.1016/j.celrep.2019.02.067 (2019).

30 Silva, G. M., Finley, D. & Vogel, C. K63 polyubiquitination is a new modulator of the oxidative stress response. Nat Struct Mol Biol 22, 116–123, doi: 10.1038/nsmb.2955 (2015).

31 Back, S., Gorman, A. W., Vogel, C. & Silva, G. M. Site-Specific K63 Ubiquitinomics Provides Insights into Translation Regulation under Stress. J Proteome Res 18, 309–318, doi: 10.1021/acs.jproteome.8b00623 (2019).

32 Higgins, R. et al. The Unfolded Protein Response Triggers Site-Specific Regulatory Ubiquitylation of 40S Ribosomal Proteins. Mol Cell, doi: 10.1016/j.molcel.2015.04.026 (2015).

33 Collart, M. A. & Panasenko, O. O. The Ccr4--not complex. Gene 492, 42–53, doi: 10.1016/j.gene.2011.09.033 (2012).

34 Panasenko, O. O. & Collart, M. A. Presence of Not5 and ubiquitinated Rps7A in polysome fractions depends upon the Not4 E3 ligase. Molecular microbiology 83, 640–653, doi: 10.1111/j.1365-2958.2011.07957.x (2012).

35 Preissler, S. et al. Not4-dependent translational repression is important for cellular protein homeostasis in yeast. EMBO J 34, 1905–1924, doi: 10.15252/embj.201490194 (2015).

36 Ikeuchi, K. et al. Collided ribosomes form a unique structural interface to induce Hel2-driven quality control pathways. The EMBO journal, doi: 10.15252/embj.2018100276 (2019).

37 Van Dalfsen, K. M. et al. Global Proteome Remodeling during ER Stress Involves Hac1-Driven Expression of Long Undecoded Transcript Isoforms. Dev Cell 46, 219–235 e218, doi: 10.1016/j.devcel.2018.06.016 (2018).

38 Absmeier, E., Santos, K. F. & Wahl, M. C. Functions and regulation of the Brr2 RNA helicase during splicing. Cell Cycle 15, 3362–3377, doi: 10.1080/15384101.2016.1249549 (2016).

39 Di Santo, R., Aboulhouda, S. & Weinberg, D. E. The fail-safe mechanism of post-transcriptional silencing of unspliced HAC1 mRNA. Elife 5, doi: 10.7554/eLife.20069 (2016).

40 Matsuki, Y. et al. Crucial role of leaky initiation of uORF3 in the downregulation of HNT1 by ER stress. Biochem Biophys Res Commun, doi: 10.1016/j.bbrc.2020.04.104 (2020).

41 Li, K., Ossareh-Nazari, B., Liu, X., Dargemont, C. & Marmorstein, R. Molecular basis for bre5 cofactor recognition by the ubp3 deubiquitylating enzyme. J Mol Biol 372, 194–204, doi: 10.1016/j.jmb.2007.06.052 (2007).

42 Muller, M. et al. Synthetic quantitative array technology identifies the Ubp3-Bre5 deubiquitinase complex as a negative regulator of mitophagy. Cell Rep 10, 1215–1225, doi: 10.1016/j.celrep.2015.01.044 (2015).

43 Matsuki, Y. et al. Crucial role of leaky initiation of uORF3 in the downregulation of HNT1 by ER stress. Biochem Biophys Res Commun 528, 186–192, doi: 10.1016/j.bbrc.2020.04.104 (2020).

44 Mohammad, M. P., Munzarova Pondelickova, V., Zeman, J., Gunisova, S. & Valasek, L. S. In vivo evidence that eIF3 stays bound to ribosomes elongating and terminating on short upstream ORFs to promote reinitiation. Nucleic Acids Res 45, 2658–2674, doi: 10.1093/nar/gkx049 (2017).

45 Munzarova, V. et al. Translation reinitiation relies on the interaction between eIF3a/TIF32 and progressively folded cis-acting mRNA elements preceding short uORFs. PLoS Genet 7, e1002137, doi: 10.1371/journal.pgen.1002137 (2011).

46 Khoshnevis, S. et al. Structural integrity of the PCI domain of eIF3a/TIF32 is required for mRNA recruitment to the 43S pre-initiation complexes. Nucleic Acids Res 42, 4123–4139, doi: 10.1093/nar/gkt1369 (2014).

47 Aitken, C. E. et al. Eukaryotic translation initiation factor 3 plays distinct roles at the mRNA entry and exit channels of the ribosomal preinitiation complex. Elife 5, doi: 10.7554/eLife.20934 (2016).

48 des Georges, A. et al. Structure of mammalian eIF3 in the context of the 43S preinitiation complex. Nature 525, 491–495, doi: 10.1038/nature14891 (2015).

49 Llácer, J. L. et al. Conformational Differences between Open and Closed States of the Eukaryotic Translation Initiation Complex. Mol Cell 59, 399–412, doi: 10.1016/j.molcel.2015.06.033 (2015).

50 Longtine, M. S. et al. Additional modules for versatile and economical PCR-based gene deletion and modification in Saccharomyces cerevisiae. Yeast 14, 953–961, doi: 10.1002/(SICI)1097-0061(199807)14:10<953::AID-YEA293>3.0.CO;2-U (1998).

51 Janke, C. et al. A versatile toolbox for PCR-based tagging of yeast genes: new fluorescent proteins, more markers and promoter substitution cassettes. Yeast 21, 947–962, doi: 10.1002/yea.1142 (2004).

